# Prediction error neurons in mouse cortex are molecularly targetable cell types

**DOI:** 10.1101/2022.07.20.500837

**Authors:** Sean M. O’Toole, Hassana K. Oyibo, Georg B. Keller

## Abstract

Predictive processing postulates the existence of prediction error neurons in cortex. Functionally, both negative and positive prediction error neurons have been identified in layer 2/3 of visual cortex, but whether they correspond to transcriptionally defined subpopulations is unclear. Here we used the activity-dependent, photoconvertible marker CaMPARI2 to tag neurons in layer 2/3 of visual cortex during stimuli and behaviors designed to trigger prediction errors. We performed single-cell RNA-sequencing on these populations and found that previously annotated *Adamts2* and *Rrad* layer 2/3 cell types were enriched when photolabeling for negative or positive prediction error responses respectively. Finally, we validated these results functionally by designing artificial promoters for use in AAV vectors to express genetically encoded calcium indicators. Thus, positive and negative prediction error responses mapped onto transcriptionally distinct cell types in layer 2/3 that can be targeted using AAV vectors.

## INTRODUCTION

Predictive processing is a computational framework that has been proposed to explain cortical function (Bastos et al., 2012; Clark, 2013; Jordan and Rumelhart, 1992; Keller and Mrsic-Flogel, 2018; Koster-Hale and Saxe, 2013; Rao and Ballard, 1999) as well as the symptoms of cortical dysfunction (Griffin and Fletcher, 2017; Palmer et al., 2015). It postulates that cortex maintains an internal representation of the external world by integrating over prediction errors computed between sensory input and predictions of that input. Predictions originate in other cortical areas and are exchanged via long range cortico-cortical projections (Garner and Keller, 2022; Leinweber et al., 2017). Prediction errors are thought to be computed in a dedicated population of neurons referred to as prediction error neurons. These come in two variants that are referred to as positive and negative prediction error neurons. A positive prediction error signals the unpredicted appearance of a sensory stimulus, and a negative prediction error signals an absence or cessation of a predicted sensory stimulus. While the computational ideas of predictive processing are well established, physiological evidence for the different cell types postulated to exist in cortex began to emerge only in the past decade. A central tenet in the experimental approach to testing the framework has been the fact that one of the best predictors of sensory input is self-generated movement. Almost all change to visual input is the direct consequence of self-generated eye, head, or body movements. Using movement as an experimental proxy for movement-related predictions, a mismatch between movement and resultant sensory feedback can be used to probe for sensorimotor prediction errors. In sensory cortices, sensorimotor prediction error responses were first described in human primary visual cortex (V1) (Stanley and Miall, 2007), and later also found across a range of species and cortical areas (Audette et al., 2021; Ayaz et al., 2019; Eliades and Wang, 2008; Heindorf et al., 2018; Keller and Hahnloser, 2009). At the same time, evidence began to emerge that in layer 2/3 (L2/3) of cortex, positive and negative prediction error neurons constitute functionally distinct sets of neurons (Attinger et al., 2017; Audette et al., 2021; Jordan and Keller, 2020; Keller et al., 2012; Zmarz and Keller, 2016). PPE neurons are thought to be excited by sensory input and inhibited by movement related predictions, while NPE neurons are driven by movement related predictions and inhibited by sensory input.

In L2/3 of mouse V1 three types of excitatory neurons have been identified, each named after one of the differentially expressed genes: *Adamts2, Agmat*, and *Rrad* (Tasic et al., 2018). These L2/3 excitatory cell types correspond to layer 2 (L2) and layer 3 (L3) transcriptional classes of excitatory neurons found in human cortex (Hodge et al., 2019). In somatosensory cortex, it has been demonstrated that the L2/3 *Rrad* type is largely driven by feedforward sensory input (Condylis et al., 2022), and in inhibitory neurons it has recently been found that a specific transcriptomic axis predicts how a neuron is functionally modulated by locomotion (Bugeon et al., 2022). Thus, we speculated that positive and negative prediction error neurons can be mapped onto transcriptionally defined neuron types within L2/3 of V1. Experimental and translational progress will hinge on determining whether these computationally identified neuron types can be selectively targeted for transgene expression; whether this is the case is still unclear.

## RESULTS

Our strategy to search for transcriptional markers of prediction error neurons in L2/3 of V1 was to first tag neurons by photoconversion that exhibit prediction error responses with a fluorescent label in vivo dissociate the tissue, sort the cells by fluorescence, and single-cell sequence different photoconversion groups separately (**Figure 1A**). To do this, we used CaMPARI2, an engineered fluorescent protein that can be photoconverted from green to red by illumination with violet light only when intracellular calcium levels are high (Moeyaert et al., 2018), which has been demonstrated to distinguish neurons of differing activity levels in vivo (Trojanowski et al., 2021). By timing photoconversion light to coincide with a particular stimulus, one can photo-tag neurons that exhibit calcium responses to that stimulus. All mice first received injections of AAV2/1-hSyn-NES-his-CaMPARI2-WPRE and a cranial window implant bilaterally in V1. Once CaMPARI2 was expressed (21 days after injection), all mice underwent the same visuomotor paradigm in a virtual reality environment. Mice were head-fixed on a spherical treadmill surrounded by a toroidal screen (**Figure 1A**). Over the course of 90 minutes, mice first experienced a closed-loop session in which locomotion was coupled to visual flow feedback presented on the toroidal screen. During this session, to evoke negative prediction error signals, we introduced visuomotor mismatches in the form of brief (1 s) halts of visual flow at random times. The closed-loop session was followed by a dark session. Finally, to evoke positive prediction error prediction error signals, we presented full-field drifting gratings with randomized onsets, durations, and orientations. Throughout all sessions, mice were free to run on the treadmill. In the first group of mice, we directed a violet (405 nm) laser at V1 for 1 s coincident with visuomotor mismatches. In the second group of mice, the laser was directed at V1 triggered by running onsets in the dark session. In the third group of mice, the laser was directed at V1 coincident with the onset of randomized (in orientation, direction, duration, and interstimulus interval) grating presentations when mice were stationary (**Figure 1B**).

**Figure 1.**
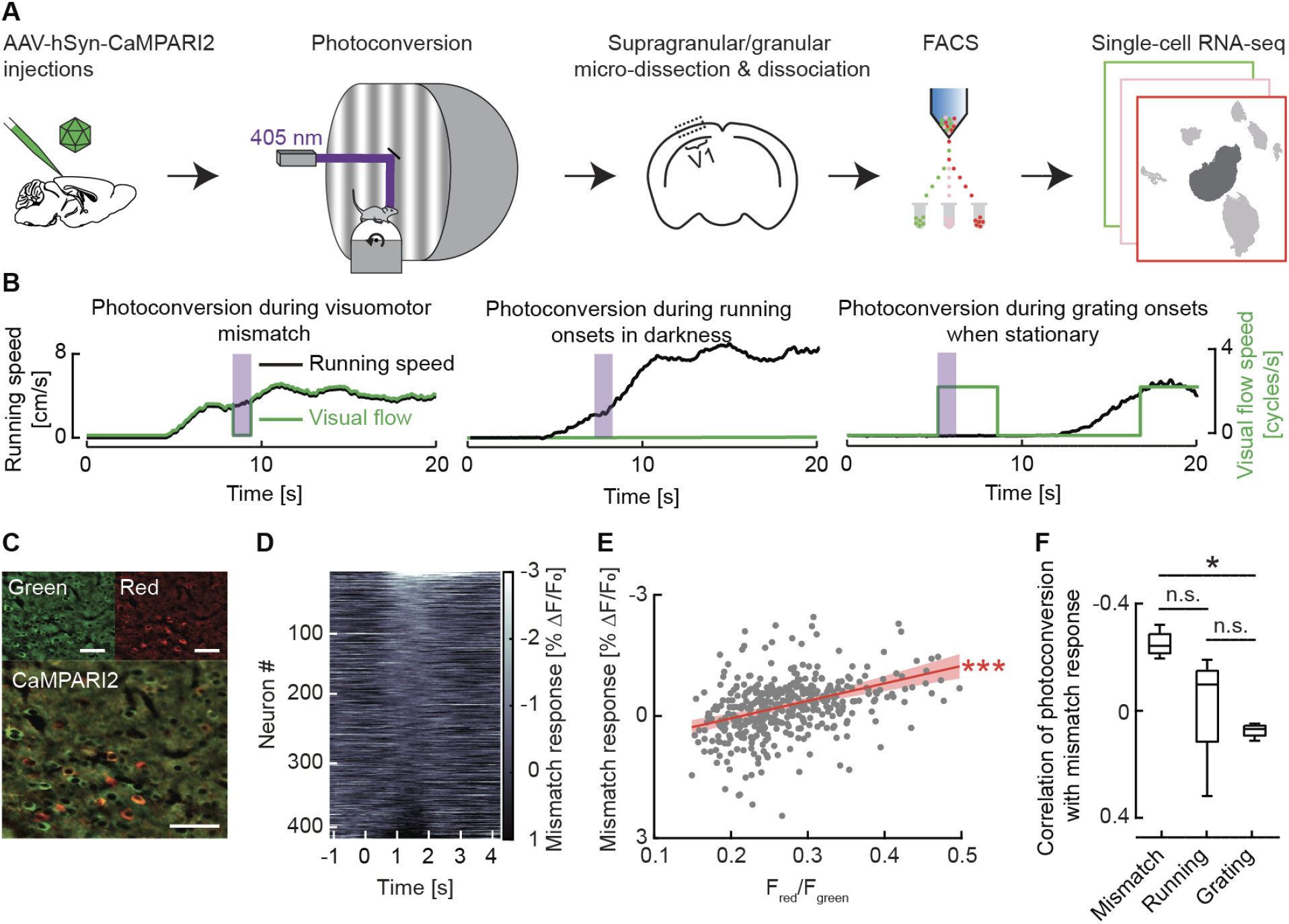
Tagging of functionally identified neurons using CaMPARI2. (**A**) Schematic of approach: C57BL/6 mice that express CaMAPRI2 in V1 were head-fixed on a spherical treadmill in a virtual reality environment. A 405 nm laser was directed at V1 through a cranial window to trigger photoconversion of CaMPARI2. Tissue from granular and supragranular V1 was dissociated, sorted into different photoconversion groups using FACS (low photoconversion is green, intermediate photoconversion is pink, high photoconversion is red), and single-cell sequenced with the 10x genomics platform. (**B**) Three separate groups of mice were used. In the first group (left), photoconversion (purple shading) was triggered on visuomotor mismatch events. In the second group of mice (middle), photoconversion was triggered on running onsets in darkness. In the third group of mice (right), photoconversion was triggered on onsets of full-field drifting grating stimuli while the mouse was stationary. (**C**) After photoconversion during visuomotor mismatch, only a subset of neurons in L2/3 that express CaMPARI2 (top left panel) were positive for the red CaMPARI2 variant (top right subpanel). Bottom image is a merge of the green and red images on top. Scale bar in all panels 25 μm. (**D**) Average CaMPARI2 responses in the green channel to visuomotor mismatch for 417 neurons recorded in 4 mice, sorted by strength of visuomotor mismatch response. Note, increases in calcium result in a decrease in fluorescence. (**E**) Scatter plot of the average fluorescence response to visuomotor mismatch plotted against the level of photoconversion per neuron, in mice that underwent photoconversion during visuomotor mismatch. The red line is an estimate of a fit to the data using linear regression (r = -0.043, intercept = 0.0091), asterisks indicate the significance of the fit. Shading indicates the 95 % confidence bounds for the linear fit. Here and elsewhere, *: p < 0.05, **: p < 0.01, ***: p < 0.001, n.s.: not significant. (**F**) Correlation of the population vector of the visuomotor mismatch, running onset, and grating response with photoconversion during mismatch for all mice. When photoconverting during visuomotor mismatch, photoconversion level and calcium response during visuomotor mismatch events were inversely correlated. This was not the case for either running or grating onset responses.

This photoconversion paradigm reliably converted a subset of CaMPARI2 expressing neurons from green to red (**Figure 1C**). To validate the targeting of photoconversion to stimulus responsive neurons, we conducted a separate set of experiments in which we measured functional responses with CaMPARI2 as well as the red to green fluorescence ratio after photoconversion. The green fluorescence of CaMPARI2 decreases when it binds to calcium, and thus CaMPARI2 can also be used as a calcium indicator. We measured calcium responses in individual neurons to visuomotor mismatches (**Figure 1D**), running onsets, and grating onsets after photoconversion during visuomotor mismatch events. We compared the level of photoconversion to the calcium responses to visuomotor mismatch in the same neurons (**Figure 1E**). There was a significant negative correlation (r = -0.25; p = 0.02) between calcium response to visuomotor mismatch and photoconversion. Note, the correlation is negative because the decrease in fluorescence corresponds to an increase in calcium levels. We found no evidence of a correlation between photoconversion and calcium responses to grating (r = 0.075, p = 0.63) or running onsets (r = 0.02, p = 0.087) (**Figure 1F**). Thus, photoconversion during visuomotor mismatch preferentially tagged visuomotor mismatch responsive neurons.

To test for differential enrichment of cell types during visuomotor behaviors we photoconverted with one of the three different stimulus paradigms (**Figure 1B**). We then sacrificed the mice, removed the brain and microdissected layer 1 (L1) through layer 4 (L4) of the photoconverted regions of V1. The tissue was dissociated into a single-cell suspension and subsequently sorted, using fluorescence-activated cell sorting (FACS), into groups of either low, intermediate, or high photoconversion (**Figure S1**). All sorted cellular particles in the three photoconversion groups were then used for single-cell RNA sequencing.

Single-cell RNA-sequencing was performed using the 10x Genomics Chromium platform. In total, we sequenced 65 002 cells across all conditions. All samples were sequenced to an average read depth of 264 million reads, which yielded a median of 3376 unique molecular identifiers (UMIs) per cell (see Methods). Given the comparatively low read depth of our data, we used a bioinformatic strategy to leverage the power published reference dataset from V1 with higher read depth (Tasic et al., 2018) for cell classification. We aligned the two datasets against each other (Welch et al., 2019) and assigned group identities to all cells using a weighted nearest neighbor algorithm (see Methods; **Figure 2A**). To confirm appropriate group assignment, we quantified the expression of marker genes for different cell types in the different groups (**Figure S2**). The distribution of assigned cell type identity for layer 2/3 types largely conformed to the expression pattern of the marker genes (**Figures 2B, 2C and 2D**). We restricted all further analysis to the group of L2/3 excitatory neurons. Within this group we assigned each cell to one of the three L2/3 excitatory cell types previously described: *Adamts2, Agmat* or *Rrad* (Tasic et al., 2018). To confirm the accuracy of the cell type assignment, we examined genes that are differentially expressed between these three types and found their expression patterns to be comparable (**Figures S2D and S2E**).

**Figure 2.**
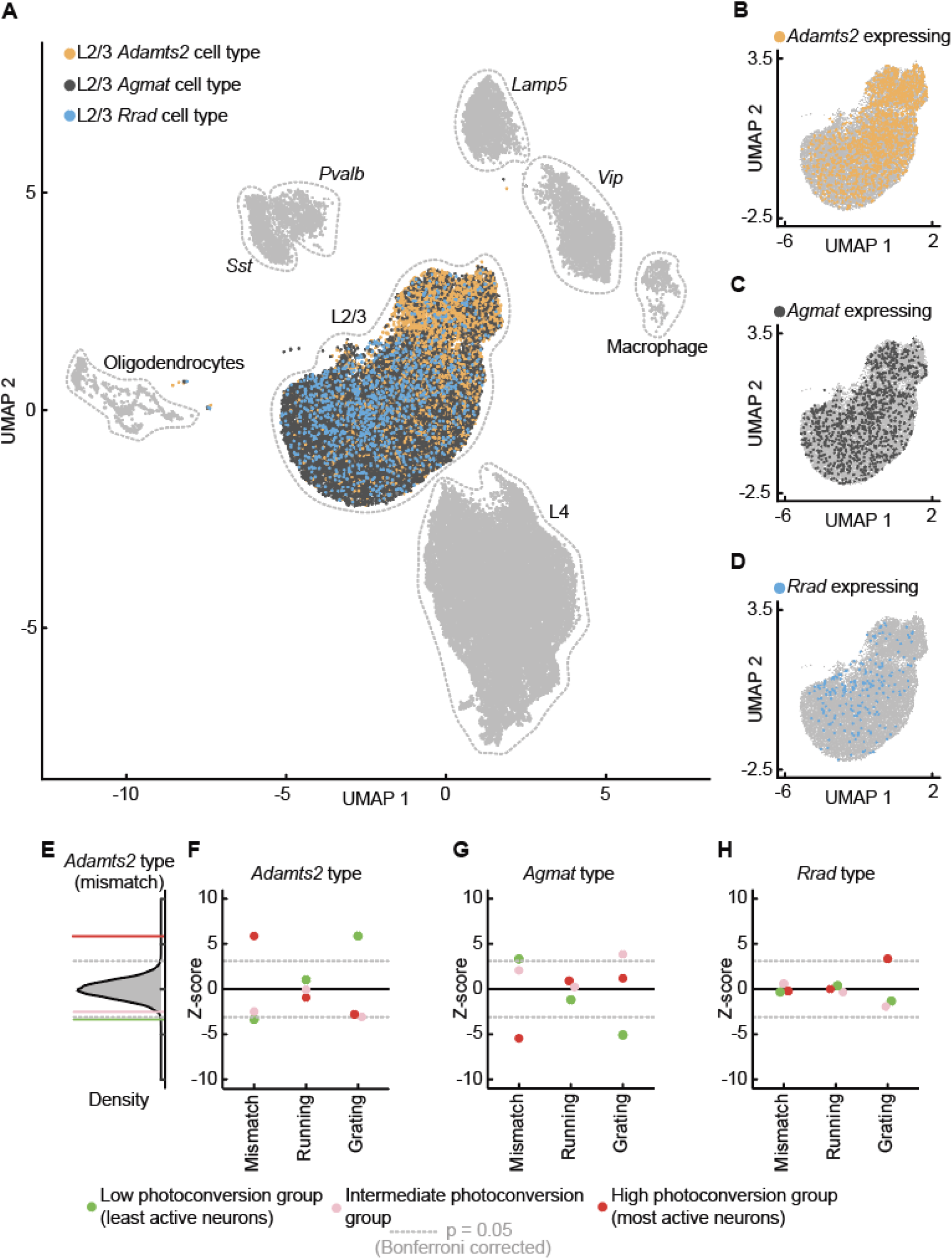
Single-cell sequencing of CaMPARI2 labelled cells revealed differential enrichment of L2/3 excitatory cell types in different visuomotor conditions. (**A**) Uniform manifold approximation and projection (UMAP) of neurons isolated from the superficial cortical layers (L1, L2/3 and L4), based on the iNMF vectors calculated from a shared projection of our data and a reference dataset (Tasic et al., 2018). Major cell groups are labelled, and inferred cell type identity of L2/3 cells are color coded. (**B**) The L2/3 cluster with cells expressing one or more UMIs corresponding to *Adamts2* highlighted. (**C**) As in **B**, but for *Agmat*. (**D**) As in **B**, but for *Rrad*. (E) Enrichment of the L2/3 *Adamts2* cell type in the high (purple), intermediate (pink), and low (green) photoconversion groups for the visuomotor mismatch labelling experiments, relative to the distribution observed in random samples (z-score distribution shown in black, dashed gray lines are the Bonferroni corrected p=0.05 levels). The L2/3 *Adamts2* cell type is significantly enriched in the high mismatch visuomotor photoconversion group. (**F**) Enrichment for the L2/3 *Adamts2* cell type across photoconversion and labelling conditions. Z-score values are displayed for visuomotor mismatch (mismatch; as in **E**), running onset (running), and grating onset when stationary (grating) labelling experiments. (**G)** As in **F**, but for the L2/3 *Agmat* cell type. (**H**) As in **F**, but for the L2/3 *Rrad* cell type.

We hypothesized that the L2/3 excitatory cell types might have different functional response properties. In the case of the L2/3 *Rrad* type, it has been found in barrel cortex of mice that *Baz1a* positive neurons (one of the marker genes of the *Rrad* type) are highly responsive to sensory stimuli (Condylis et al., 2022). To test whether each of these cell types had distinct functional response properties, we investigated whether there was a differential enrichment of the three L2/3 excitatory cell types (*Adamts2, Agmat* or *Rrad*) across stimulus paradigms (visuomotor mismatch, running onsets in darkness, or grating onsets) and photoconversion state (low, medium, or high). We found that the L2/3 *Adamts2* type was significantly overrepresented in the high visuomotor mismatch photoconversion group (**Figures 2E and 2F**). The *Agmat* type was underrepresented in both high visuomotor mismatch and low grating photoconversion groups (**Figure 2G**), and the L2/3 *Rrad* type was significantly overrepresented in the high grating photoconversion group (**Figure 2H**). These results suggest that on average visuomotor mismatch responsive neurons are enriched in the L2/3 *Adamts2* cell type, and that, consistent with previous findings (Condylis et al., 2022), neurons responding to feed-forward sensory stimuli are enriched in the L2/3 *Rrad* cell type. The remaining L2/3 *Agmat* type appears to represent an intermediate population or potentially a grouping with an as yet unidentified function.

To confirm the functional identity of these cell types, we designed artificial promoters (AP) for use in AAV vectors to target the expression of a genetically encoded calcium indicator to each one of these cell types using a previously described strategy (Jüttner et al., 2019). Artificial promoters were designed based on 2 kb segments surrounding the transcription start site of selected marker genes corresponding to the three cell types (**Figure 3A**). To test for expression, we first used these promoters to drive GFP expression and examined the expression profiles visually. The successful candidate for the L2/3 *Adamts2* cell type was a promoter designed off of the *Adamts2* gene (AP.Adamts2.1), for the L2/3 *Agmat* cell type, one based on the *Agmat* gene (AP.Agmat.1), and for the L2/3 *Rrad* cell type, one based on the *Baz1a* gene (AP.Baz1a.1). We initially tried labelling the L2/3 *Rrad* cell type with an AP based on the Rrad gene, however, expression was too dim and extremely sparse (**Figure S3**). Each of these three artificial promoters resulted in a different expression profile in cortex (**Figures 3B, 3C and 3D**). AP.Adamts2.1 drove expression in superficial L2/3, in superficial L5, and in L6 (**Figures 3B and 3F**). AP.Agmat.1 drove expression throughout L2/3, and exhibited very little expression in other layers (**Figures 3C and 3F**).

**Figure 3.**
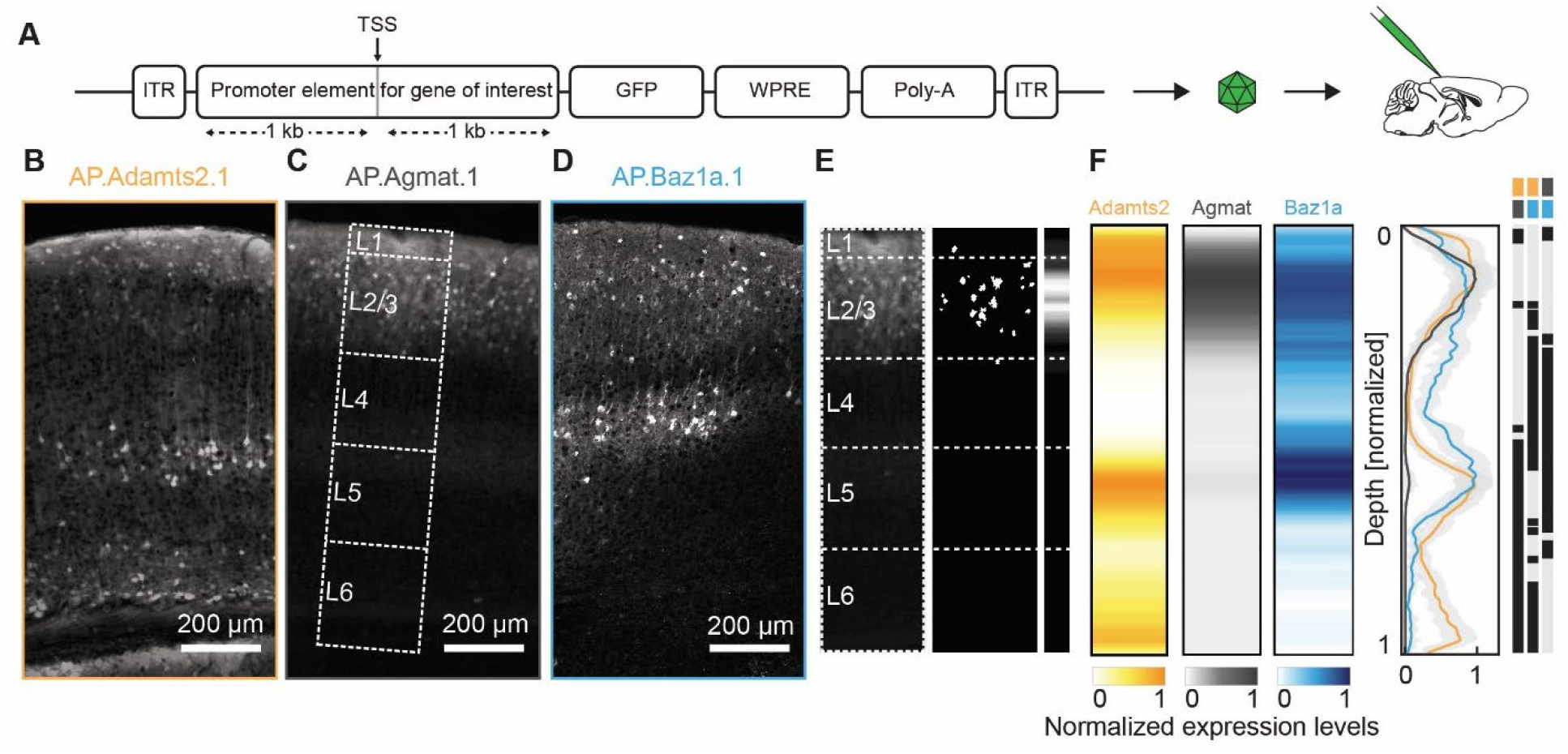
Artificial promoters exhibit differential expression patterns in cortex. (**A**) Artificial promoter design strategy. The 2 kb segment of the genomic sequence flanking the transcription start site (TSS), for the selected marker genes, was placed into an AAV vector to drive GFP expression. (**B**) Example expression pattern from an AAV2/1-AP.Adamts2.1-GFP injection in V1. (**C**) As in **B**, but for AAV2/1-AP.Agmat.1-GFP. (**D**) As in **B**, but for AAV2/1-AP.Baz1a.1-GFP. (**E**) Histology images were converted to expression profiles as a function of cortical depth (see Methods). The example section is shown as an inset in **C.** (**F**) Left: Average expression profiles as a function of cortical depth for the three different artificial promoters. Right: Comparison between the expression profiles. Significant differences are marked as black bars to the right of the plot, and the respective comparisons are labelled by the pair of line segments at the top.

AP.Baz1a.1 drove expression in L2/3, L4, and L5 (**Figures 3D and 3F**). For each of these promoters we sequenced FACS sorted cells with bulk RNA-sequencing to confirm marker gene expression in the targeted neurons (**Figure S4**). Also, for AP.Baz1a.1 and AP.Adamts2.1, we noticed a large number of neurons labelled in L1, indicating that these promoters also induce transgene expression in L1. Using RNA-sequencing and histology experiments, we confirmed that indeed a subset of the labelled cells were inhibitory neurons. To prevent expression in inhibitory neurons, we designed Cre dependent variants of the AP.Baz1a.1 and AP.Adamts2.1 vectors and used them in an Emx1-Cre mouse line to restrict expression to excitatory cortical neurons.

Using these promoters, we could then target expression of a GCaMP variant to the three L2/3 excitatory cell types separately to confirm their functional specificity. We did this in three groups of mice that received injections of either AAV2/1-AP.Adamts2.1-DIO-GcaMP8s, AAV2/1-AP.Agmat.1-GcaMP7f, or AAV2/1-AP.Baz1a.1-DIO-GcaMP8s in V1. To measure functional responses in these neurons using two-photon calcium imaging, mice underwent the same visuomotor paradigm (closed-loop, running in darkness, randomized grating presentations) as during photoconversion experiments with the exception that the mice were also exposed to an open-loop session during which the visual flow the mice had self-generated in the preceding closed-loop session was replayed. Thus, we asked whether the response profiles of these cell types behaved in a way similar to the observations found in the single-cell sequencing data analysis.

The AP.Adamts2.1 neurons responded robustly to visuomotor mismatch as well as to running onsets (in darkness, closed-loop or open-loop sessions) and decreased their fluorescence following grating onsets (**Figures 4A, 4B and 4C**). While AP.Agmat.1 neurons responded primarily to running onsets, AP.Baz1a.1 neurons exhibited a robust grating onset response. The visuomotor mismatch responses were significantly larger in the population of AP.Adamts2.1 neurons than they were in the AP.Baz1a.1 and AP.Agmat.1 populations (**Figure 4A**). Conversely, the grating onset response was stronger in the AP.Baz1a.1 population than in the other two labelled populations (**Figure 4C**). Consistent with an opposing influence of visual flow and motor related input on negative prediction error and positive prediction error neurons (Attinger et al., 2017; Jordan and Keller, 2020), we found that the activity of AP.Baz1a.1 neurons was positively correlated with visual flow speed, while that of AP.Adamts2.1 neurons was negatively correlated (**Figure 4D**). For correlations with running speed, this was reversed in that AP.Adamts2.1 neurons exhibited the strongest correlations, and AP.Baz1a.1 neurons the weakest correlations (**Figure 4E**). The activity of AP.Agmat.1 neurons was uncorrelated with visual flow speed, and exhibited an intermediate correlation with running speed.

**Figure 4.**
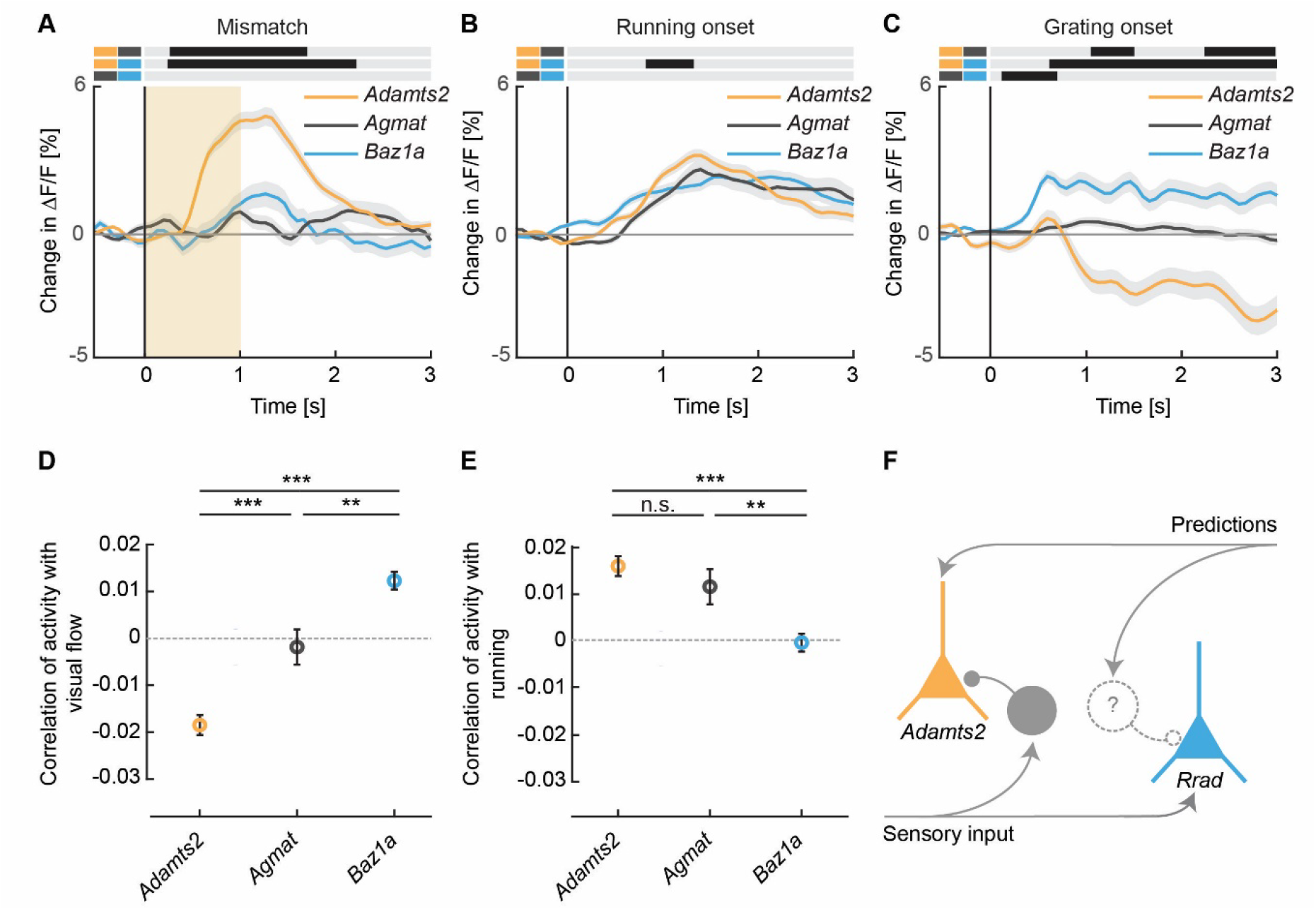
L2/3 neurons targeted by artificial promoters exhibited differential visuomotor responses. (**A**) Average response to visuomotor mismatch in the AP.Adamts2.1 labelled population (orange) was stronger than in the AP.Agmat.1 (dark gray), or the AP.Baz1a.1 (blue) populations. Orange shading indicates the duration of visuomotor mismatch; grey shading indicates the SEM. Response curves were compared bin-by-bin (p < 0.05 indicated by black bars above traces). Each of the three comparisons is denoted by a pair of line segments to the left, corresponding in color to the data being plotted. (**B**) As in **A**, but for the response to running onsets. (**C**) As in **A**, but for the response to the onset of randomized grating. (**D**) Average correlation of neuronal activity with visual flow during open-loop sessions for L2/3 *Adamts2, Agmat*, and *Baz1a* neurons. Error bars are SEM over neurons. Here and elsewhere, *: p < 0.05, **: p < 0.01, ***: p < 0.001, n.s.: not significant. (**E**) As in **D**, but for average correlation of neuronal activity with running during open-loop sessions. Error bars are SEM over neurons. (**F**) Schematic of a model circuit that would explain the response properties of *Adamts2* and *Rrad* L2/3 excitatory neuron types. Triangles, indicate L2/3 excitatory neuron types; circles, indicate inhibitory interneurons. L2/3 *Adamts2* neurons respond robustly to visuomotor mismatch, and their activity is correlated with running, and negatively correlated with visual flow. This is consistent with the functional signature of a negative prediction error type. L2/3 *Rrad* neurons have visually driven responses consistent with a positive prediction error type. Previous work has demonstrated that negative prediction error neurons are bottom-up inhibited (Attinger et al., 2017), while a potential source of top-down driven inhibition for the positive prediction error neuron remains speculative.

## DISCUSSION

In our interpretation, we have made the assumption that visually responsive excitatory neurons in L2/3 are positive prediction error neurons, and that visuomotor mismatch responsive neurons are negative prediction error neurons. It should be noted, that exhibiting visual or visuomotor mismatch responses does not mean a neuron functions as a positive prediction error or negative prediction error neuron.

However, our interpretation is based on previous work demonstrating that in L2/3, visual and visuomotor mismatch responsive neurons function as comparators between top-down and bottom-up input (Jordan and Keller, 2020). Based on our findings, the L2/3 *Adamts2* population is enriched for negative prediction error neurons, and that of L2/3 *Rrad* is enriched for positive prediction error neurons. Interestingly, we find that the third type of excitatory L2/3 neuron, the L2/3 *Agmat* cell type, does not exhibit responses consistent with visuomotor prediction errors. One possibility is that this population functions to maintain an internal representation, by integrating over negative prediction error and positive prediction error responses, a function that is typically speculated to be performed by L5 neurons (Heindorf and Keller, 2022). Another possibility is that these neurons compute prediction errors that are not visuomotor. The populations of neurons that respond e.g. to spatio-visual prediction errors, is separate from that responding to visuomotor prediction errors (Fiser et al., 2016). Lastly, it is possible that these neurons perform a function outside of the canonical circuit for predictive processing. Identifying how the three different types of L2/3 excitatory neurons interact with each other and the rest of the cortical circuit, will likely be critical for making progress in our understanding of the canonical cortical circuit.

Our results demonstrate that prediction error responsive neurons have distinct transcriptional identities and can be targeted virally. Given that predictive processing has become one of the leading theories of cortical function (Bastos et al., 2012; Clark, 2013; Jordan and Rumelhart, 1992; Keller and Mrsic-Flogel, 2018; Koster-Hale and Saxe, 2013; Rao and Ballard, 1999) with the capacity to explain both neurophysiological results, as well as dysfunctions of cortex that are thought to result in diseases like autism (Lawson et al., 2014, 2017; Sinha et al., 2014) and schizophrenia (Corlett et al., 2009, 2009; Fletcher and Frith, 2009; Frith et al., 2000), it will be essential to have specific genetic access to the functionally identified neuronal subpopulations postulated by this framework. Experimental tests of different variants of predictive processing circuit models will hinge on cell type specific manipulations and recordings of neuronal activity (Keller and Mrsic-Flogel, 2018). In parallel, it is likely that any treatment of disorders involving prediction error computations, will require cell type specific interventions. The identification of *Adamts2* as a marker of neurons with negative prediction error responses and that of *Baz1a* as a marker of neurons with positive prediction error responses in L2/3 of mouse cortex is a first step in this direction.

## SUPPLEMENTARY FIGURES

**Figure S1.**
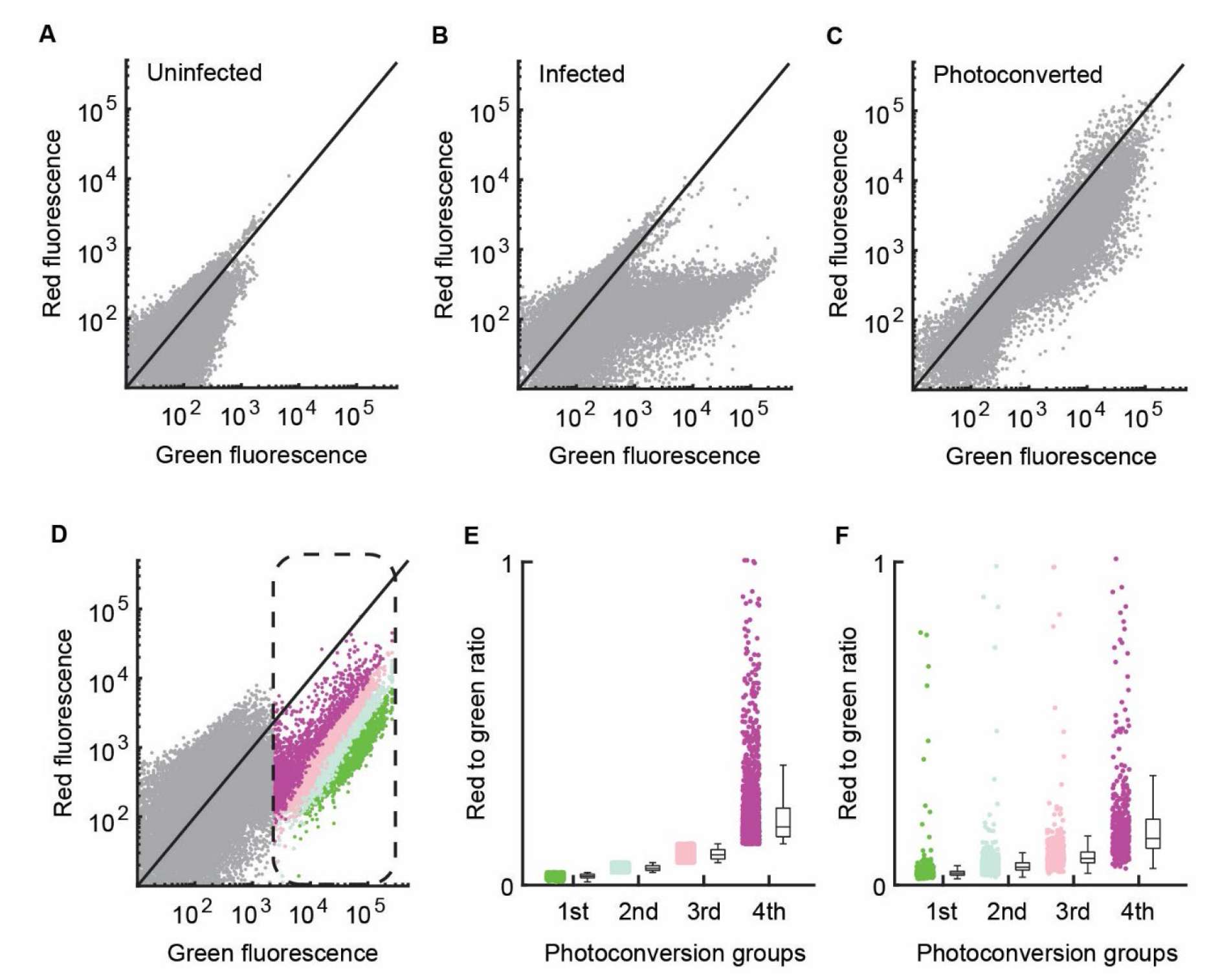
FACS sorting of CaMPARI2 infected cells that underwent photoconversion. (**A**) FACS plot of an example dissociated piece of cortical tissue without CaMPARI2 expression. Each dot corresponds to one putative cell. Cells were selected for size, filtered for doublets, and high Draq7 (dead) cells were excluded. (**B**) As in **A**, but for an infected cortical preparation that expressed CaMPARI2 without photoconversion. (**C**) As in **B**, with photoconversion. (**D**) An example FACS plot illustrating ratio-metric sorting. FACS gates were set to exclude background fluorescence and to sort by red to green ratio. Cells were sorted into equally sized bins corresponding to the first (green), second (pale blue), third and fourth quantiles of the red to green fluorescence ratio distribution. The middle quartiles were later combined for single-cell analysis. (**E**) The distribution of red to green ratio values for cells shown in **D**, for the different photoconversion groups separately. A small number of cells had ratio values larger than 1 (not shown). (**F**) To test for stability of the FACS sorting, we measured the red to green ratio in the photoconversion groups shown in **E** separately by passing them through the FACS sorter again. Shown is the distribution of red to green ratios measured in the second measurement.

**Figure S2.**
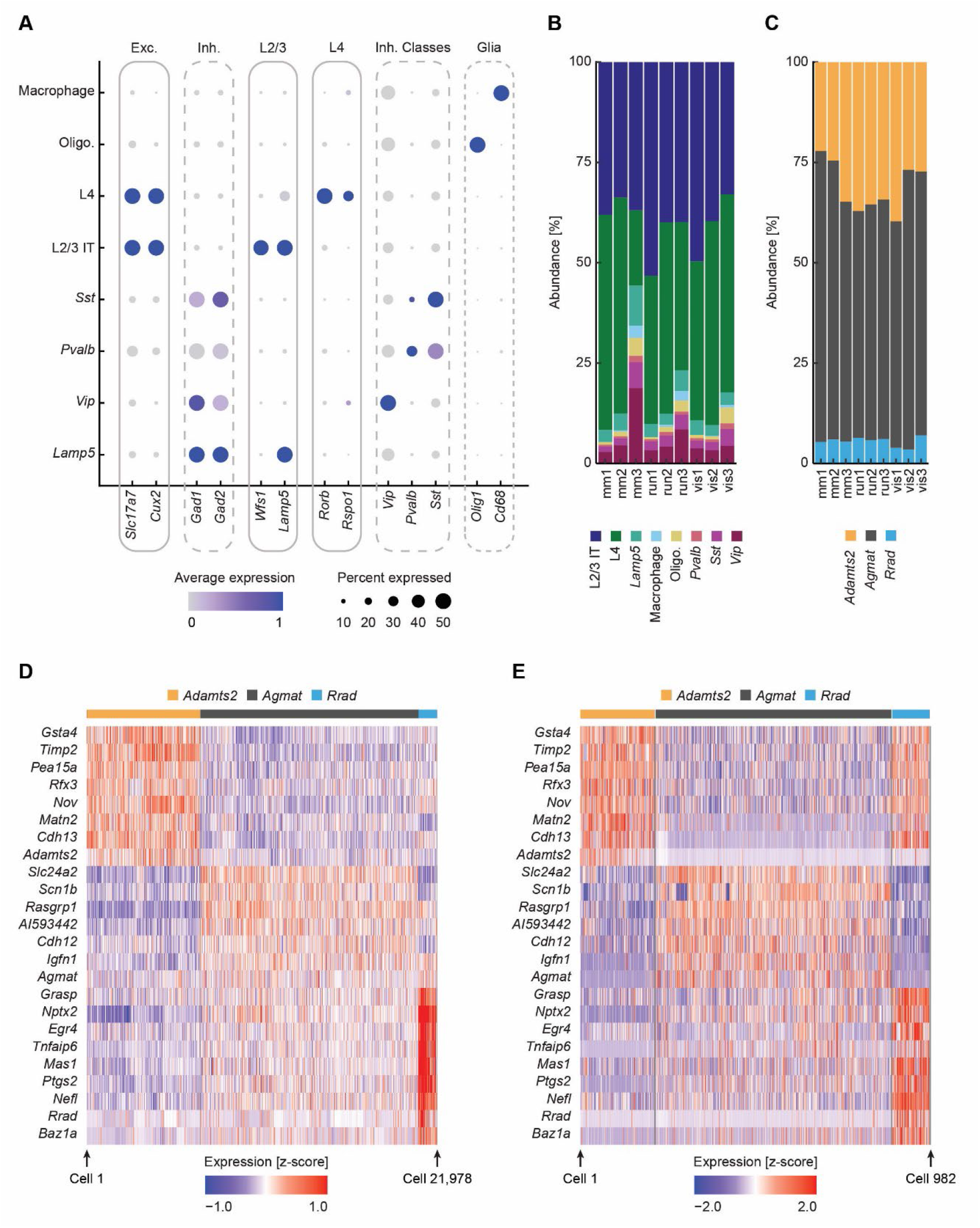
Cell type assignment accuracy. (**A**) Dot plot for a selected number of marker genes for each of the major cell groups (with a cumulative population size greater than 500 cells across all samples). Dot size represents the percentage of cells (**B**) expressing each marker while the color represents the average normalized expression value. Exc.: excitatory neurons, Inh.: inhibitory neurons, Oligo.: Oligodendrocytes, IT: intertelencephalic. (**C**) Abundance of each major cell group as shown in **A**, across the three photoconversion groups and the three photoconversion types. mm: Mismatch; run: Running onsets; vis: Grating onsets; 1: Low photoconversion group; 2: Intermediate photoconversion group; 3: High photoconversion group. (**D**) As in **B**, but for the three L2/3 excitatory types. (**E**) Heatmap for a select number of genes enriched in each of the three L2/3 excitatory cell types. There were 21 978 L2/3 excitatory neurons examined in this study (7113 *Adamts2*, 13 688 *Agmat*, 1177 *Rrad*, and 64 unassigned neurons, the latter are not shown in the heatmap). Color values correspond to expression levels measured as a z-score. To prevent aliasing, data are smoothed across cells with a window size of 20. (**F**) As in **D**, for all L2/3 V1 excitatory neurons from a reference dataset (Tasic et al., 2018).

**Figure S3.**
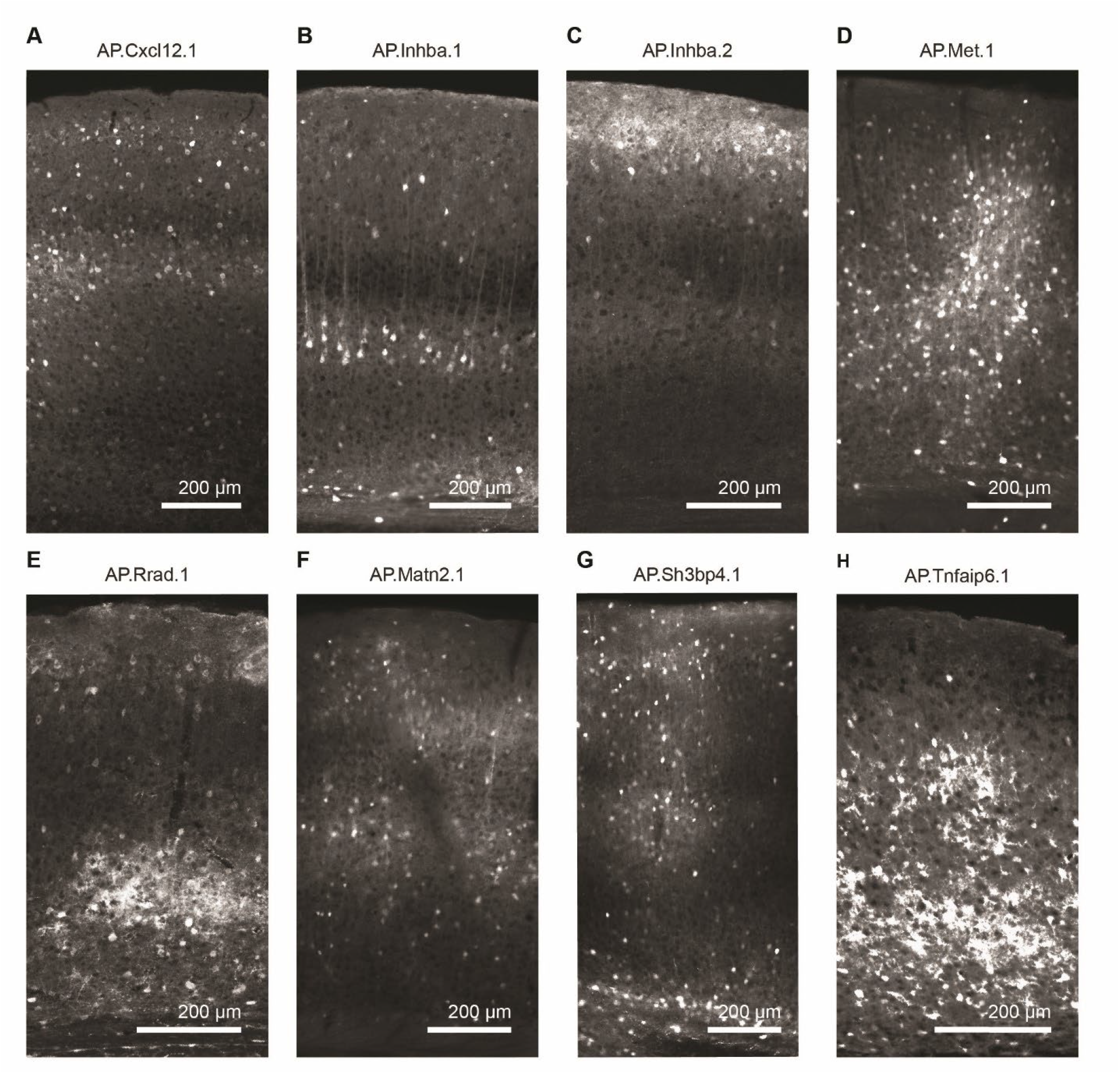
Some of the failed attempts of promoter designs. (**A**) Expression pattern of an AAV2/1-AP.Cxcl2.1-GFP with an artificial promoter designed based on the 2 kb surrounding the transcription start site of *Cxcl12*, a putative maker of the L2/3 *Agmat* cell type. (**B**) As in **A**, but for an artificial promoter designed based on the 2 kb surrounding the first transcription start site (relative to the + strand of chromosome 13) of the *Inhba* gene, a putative marker of the L2/3 *Adamts2* cell type. (**C**) As in **A**, but for an artificial promoter designed based on the 2 kb surrounding the second transcription start site (relative to the + strand of chromosome 13) of the *Inhba* gene, a putative marker of the L2/3 *Adamts2* cell type. (**D**) As in **A**, but for an artificial promoter designed based on the 2 kb surrounding the first transcription start site (relative to the + strand of chromosome 6) of the *Met* gene, a putative marker of the L2/3 *Adamts2* cell type. (**E**) As in **A**, but for an artificial promoter designed based on the 2 kb surrounding the transcription start site of the *Rrad* gene, a putative marker of the L2/3 *Rrad* cell type. (**F**) As in **A**, but for an artificial promoter designed based on the 2 kb surrounding the transcription start site of the *Matn2* gene, a putative marker of the L2/3 *Adamts2* cell type. (**G**) As in **A**, but for an artificial promoter designed based on the 2 kb surrounding the transcription start site of the *Sh3bp4* gene, putative marker of the L2/3 *Adamts2* cell type. (**H**) As in **A**, but for an artificial promoter designed based on the 2 kb surrounding the transcription start site of the *Tnfaip6* gene, putative marker of the L2/3 *Rrad* cell type.

**Figure S4.**
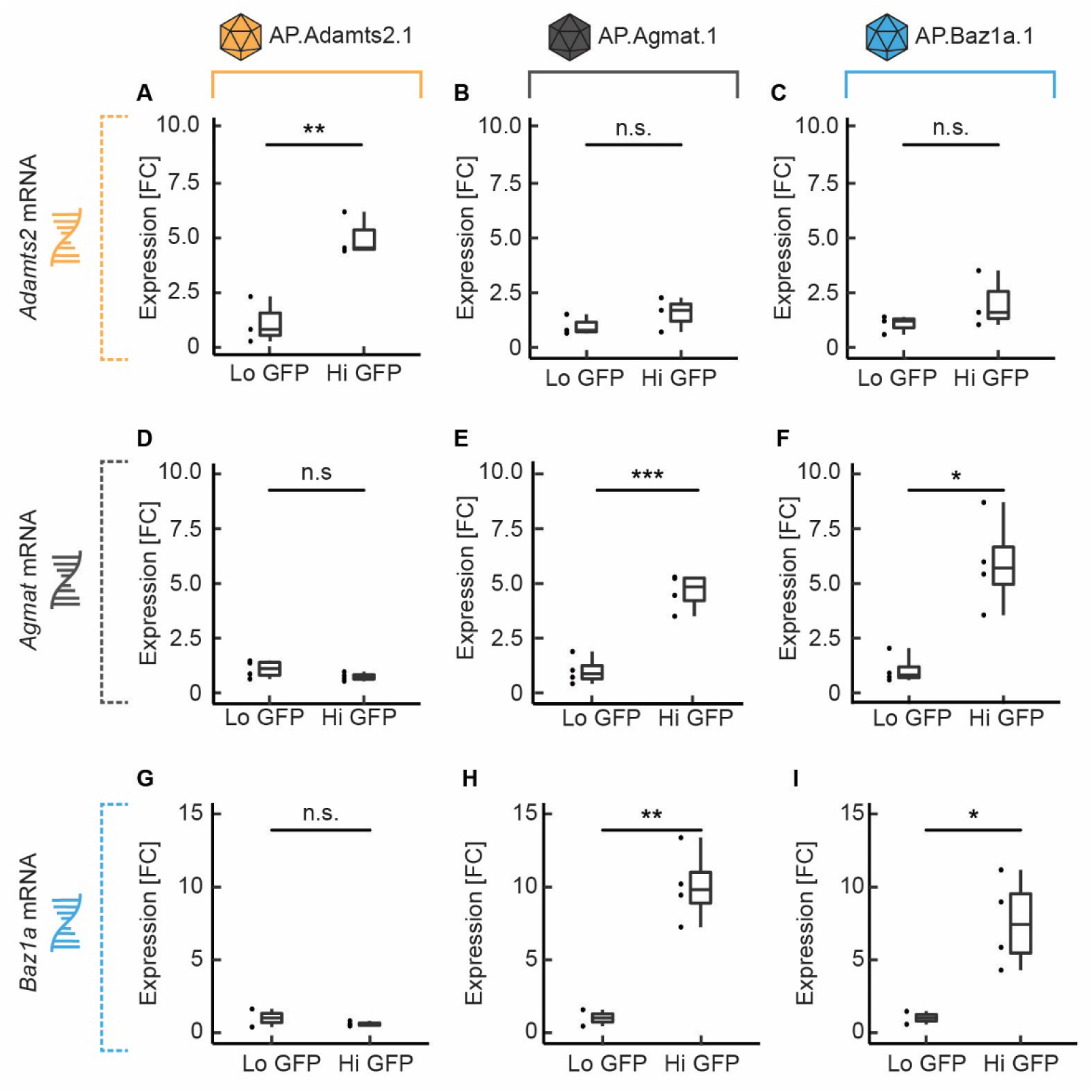
Differential expression of marker genes for artificial promoter infected neurons. (**A**) Bulk RNA-sequencing data for *Adamts2* expression in populations of L1, L2/3 and L4 cortical neurons infected with an AAV2/1-AP.Adamts2.1-GFP, for high (Hi) and low (Lo) GFP expression level populations separately. The low and high GFP groups constitute the lower and upper thirds of the GFP fluorescence distribution (cells with no expression were excluded). Fold-change (FC) values were normalized to the average expression in the low GFP group. Note all samples were collected in pairs, however, in some cases library preparation failed for one of the two samples thus sample sizes were unequal (here and in other figure panels of this figure). (**B**) As in **A**, except for the AP.Agmat.1 promoter. (**C**) As in **A**, except for the AP.Baz1a.1 promoter. (**D-F**) As in **A-C**, except for *Agmat* expression. (**G-I**) As in **A-C**, except for *Baz1a* expression.

## MATERIALS AND METHODS

### Mice and surgical procedures

All animal procedures were approved by and carried out in accordance with guidelines of the Veterinary Department of the Canton Basel-Stadt, Switzerland. For most experiments we used C57BL/6J mice (Charles River). For the experiments designed to restrict transgene expression to excitatory neurons we used Emx-Cre knock-in mice (Gorski et al., 2002). For all surgical procedures, mice were anesthetized with a mixture of Fentanyl (0.05 mg/kg; Actavis), Midazolam (5.0 mg/kg; Dormicum, 603 Roche) and Medetomidine (0.5 mg/kg; Domitor, Orion). In all mice used for functional imaging, a 4 mm or 5 mm craniotomy was made over right V1 and 4 injections of approximately 250 nl each were made centered on 2.5 mm lateral and 0.5 mm anterior of lambda. For CaMPARI2 single-cell sequencing experiments two 4 mm windows were implanted at the same coordinates bilaterally with two injections in each hemisphere. Circular glass cover slips were glued (Ultragel, Pattex) in place to close the craniotomy, and the exposed skull covered in Histoacryl (Braun). To enable head-fixation, a custom-made titanium head-plate was attached to the skull with dental cement (Heraeus Kulzer).

### Acclimatization to experimental setups

Prior to the start of photoconversion and imaging experiments, mice were trained on the experimental setups for 3 days in the case of photoconversion experiments, and 6 days for functional imaging experiments. Each training session lasted for 1 hour during which the mouse was head-fixed on the spherical treadmill and experienced closed-loop visual flow feedback projected onto a toroidal screen surrounding the mouse. The rotation of the treadmill was restricted to rotation around the transverse axis.

### Photoconversion

Photoconversion experiments were performed on mice that robustly expressed CaMPARI2 throughout left and right V1. Out of a total of 34 mice, 12 (4 for each condition) were selected for photoconversion experiments based on CaMPARI2 expression levels, vasculature, absence of any signs of infection or noticeable bone regrowth. For photoconversion experiments, mice were head-fixed on a spherical treadmill surrounded by a toroidal screen. The screen covered an area corresponding to 240 degrees horizontally and 100 degrees vertically relative to the mouse’s field of view. All mice were exposed to the same sequence of visuomotor sessions. First, mice experienced 30 min of closed-loop coupling between running and visual flow feedback in a virtual corridor. To induce visuomotor mismatches, we briefly (1 s) halted the visual flow at random intervals (on average every 15 s, +/-10 s). Next, mice experienced 30 min of a dark session during which all light sources were either turned off or shielded. Finally, mice were presented with full-field drifting sinusoidal grating stimuli (2.2 Hz, 0.04 cycles per degree), designed to be minimally predictable, for 3 to 8 s of randomly selected directions (0°, 45°, 90°, 135°, 180°, 225°, or 270°), spaced by 2 s to 6 s gray screen presentations, for a total of 30 min. Mice were free to run throughout all sessions.

For photoconversion, we used a 405 nm laser (Obis, Coherent). The laser was directed bilaterally at V1 through the cranial windows for a duration of 1 s for each trigger. The laser power was 100 mW with a diameter of 3 mm (FWHM) at the surface of the brain, similar to that used in a previous study (Moeyaert et al., 2018). The laser was directed between a blank position and the cranial windows using a two axis galvo scan head. During the 1 s stimulation, the beam was alternated between left and right V1 at a frequency of 60 Hz. To minimize visible changes in ambient light, we shielded the entire light path up to the head-bar of the mouse. All mice received between 30 and 50 photostimulation pulses of 1 s during the 30 min photoconversion session. In instances in which mice did not receive 50 pulses, the photoconversion session was extended until the mouse received 50 pulses. In instances in which mice received 50 pulses before the end of the session, stimulation was stopped for that session. For the visuomotor mismatch photoconversion events, we triggered the laser only on mismatch events during which the running speed of the mouse was above 2 cm/s. For photoconversion during running onsets in darkness, running onsets were detected in real-time as events with increases in running speed of at least 1.4 cm/s over a span of 500 ms, and a running speed in the window -1500 ms to -500 ms before the event that did not exceed 1 cm/s. For photoconversion during grating onsets, the laser was triggered at the grating onset if the running speed of the mouse was below 0.5 cm/s. Following photoconversion experiments, all mice were immediately anesthetized, examined on an epifluorescence microscope for evidence of successful photoconversion and sacrificed for single-cell RNA-sequencing.

Due to sample processing limitations, only two session types were collected at a time. In order to balance potential batch effects, each session type was collected twice in tandem with one of the other session types. Thus, we executed the single-cell sequencing photoconversion experiment three times (mismatch with visual; mismatch with running; running with visual). So, for all photoconversion experiments, mice were split into 3 groups of 2. In the first group of 2 mice, photoconversion was triggered concurrent with visuomotor mismatch stimuli in the closed-loop session. In the second group of 2 mice, photoconversion was triggered at the onset of running in the dark session. In the third group of 2 mice, photoconversion was triggered at the onset of grating stimuli in instances when the mice were stationary. After each experiment photoconversion was confirmed by examining the cranial window with an epifluorescence microscope. This was repeated twice for a total of 12 animals.

### Solutions

We used artificial cerebrospinal fluid (ACSF) for the generation of single-cell suspensions as previously described (Hempel et al., 2007; Tasic et al., 2018), with some modifications. We used *N*-methyl-d-glucamine (NMDG) based ACSF to improve cell viability (Ting et al., 2014). The NMDG ACSF formulation, based on a previous publication (Tasic et al., 2018), consisted of NMDG (96 mM), HCl (96 mM), NaHCO_3_ (25 mM), glucose (25 mM), HEPES (20 mM), N-acetylcysteine (12 mM), MgSO_4_ (10 mM), sodium l-ascorbate (5 mM), myo-inositol (3 mM), sodium pyruvate (3 mM), KCl (2.5 mM), thiourea (2 mM), NaH_2_PO_4_ (1.25 mM), CaCl_2_ (0.5 mM), and taurine (0.01 mM). All ACSF stocks were kept at pH 7.35 and 315 mOsm. Additionally, all solutions were supplemented with 13.2 mM trehalose to enhance cell survival during the dissociation and sorting (Saxena et al., 2012). Solutions were supplemented with a transcriptional blocker, actinomycin-d (1 μg/ml, Sigma, cat. no. A1410), to reduce dissociation related expression changes (Wu et al., 2017). In addition, the ion channel blockers TTX (0.1 μM, Tocris,), DNQX (200 μM, Merck/Sigma), and APV (50 μM, Tocris) were added to prevent any activity related changes and reduce excitotoxicity (Hempel et al., 2007). The ACSF used for slicing and cooling the brain was pre-chilled to an icy slush, and bubbled with carbogen gas (95 % O_2_, 5 % CO_2_) throughout the experiment.

### Generation of single-cell suspensions

The procedure to generate a single-cell suspension was identical for both single-cell and bulk RNA-sequencing experiments. Brain extraction and cellular dissociation for all experiments followed a standard protocol (Hempel et al., 2007) with some modifications. Mice were anesthetized and decapitated. To remove the brain, the skull was cut medio-laterally directly anterior of the cerebellum, along the lateral edge of the dorsal skull, and medio-laterally above the olfactory bulb. The bone flap was then opened, and the brain removed with a spatula. With a scalpel, the brain was cut, coronally, at 1 mm anterior of bregma to form a flat surface, the anterior most portion of the brain was discarded. On this surface, the brain was then glued to a cutting platform next to an agarose block (or embedded in low-melt agarose for single-cell sequencing experiments) and rapidly immersed in ice-cold carbogenated ACSF. Brain sections (300 μm) were cut on a vibratome (7000smz-2, Campden Instruments) with a ceramic blade at an oscillation frequency of 80 Hz and a cutting speed of 200 μm/s. Sections within and around V1 were collected with a plastic Pasteur pipette and placed into a bubbled chamber filled with room-temperature ACSF and allowed to recover for at least 15 min. Subsequently, samples were placed into 30 ml of ACSF with 1 mg/ml pronase (Protease Type XIV, Sigma), along with 1 mg collagenase (Collagenase from Clostridium histolyticum, Sigma), as previously described (O’Toole et al., 2017) and digested for 75 min. Then slices were placed in ACSF (without protease/collagenase) and allowed to recover for 15 min. After that, the slices were transferred to a 30 cm petri dish containing cold carbogenated ACSF supplemented with 1 % Fetal Bovine Serum (FBS). Supragranular layers of V1 were microdissected under an epifluorescent microscope. Microdissected V1 pieces were then transferred to 1.5 ml RNase free tubes. These tissue pieces were triturated with a series of fire-polished glass Pasteur pipettes, with inner diameters of 600 μm, 300 μm, and 150 μm, that had been coated in ACSF with 1% FBS (Hempel et al., 2007). Single-cell suspensions were then strained through nylon filters (Celltrics filters, sysmex) and placed into tubes (12 × 75 mm Round Bottom Polystyrene Test Tubes, Fisher) along with 2 µl of live/dead cell stain (DRAQ7, BioLegend) per ml and then FACS sorted.

### FACS

For FACS sorting, we used either a BD FACSAria III cell sorter or a Sony MA900 cell sorter. To enrich for cellular particles, we used a series of gates based on particle size and complexity, excluding doublets and particles that stained positive for DRAQ7, a marker of dead cells. For ratiometric FACS, we ran a series of control experiments to determine the range in which GFP and RFP expression could be observed. An uninfected control was used to determine the cut off gate for viral expression (**Figure S1**). In some cases, cell samples were examined under a fluorescence microscope to check for fluorescence. At the beginning of each experiment, we sorted at least 1000 cells above the expression threshold to estimate the fluorescence gates that would bin approximately 25% of the neurons into one of four photoconversion groups (the middle two were later combined for analysis purposes), based on their red to green fluorescence ratio. Using these gates, we then sorted 10 000 neurons per photoconversion group. For each photoconversion experiment, the single cell suspensions obtained from two mice were sorted (serially) to reach a threshold of 40 000 cells. Between mice, gates were recalibrated to match the photoconversion distribution of the current sample. We confirmed sorting reliability by resorting a sorted sample and found that the difference between photoconversion groups were maintained (**Figure S1F**). All samples were sorted directly into a PBS/BSA solution such that the final concentration was 0.04 % BSA. Immediately after sorting, samples were used for Gel Bead in Emulsions and cDNA generation.

For FACS sorting of the bulk RNA-sequencing experiments we followed a similar protocol. Prior to the experiment, mice were injected at 6 to 8 weeks of age, and brains were extracted 4 weeks later to generate single-cell suspensions. We used a series of gates based on particle size, excluding doublets and particles that stained positive for DRAQ7. As before, a pilot experiment was used to determine a cutoff value for the absence of viral expression by examining an uninfected sample. Then, for infected samples, 1000 cellular particles were measured to determine the distribution of fluorescence values. Gates were established that captured four categories of fluorescence: particles with no expression as well as low, medium, and high expressing particles each corresponding to a third of the distribution of particles above the baseline cutoff for viral expression. We restricted analysis to cells expressing low (just above the expression threshold) and high amounts of GFP. These dynamically drawn gates were used to reduce batch effects caused by infection efficiency and differences in viral titer. 1000 cell particles were sorted for each group directly into lysis buffer (Norgen). Following sorting, samples were either stored at -80°C or immediately used for RNA purification.

### Sequencing library preparation

RNA for bulk RNA-sequencing samples was purified with the Single Cell RNA Purification Kit (Norgen), and columns were treated with a DNase to remove genomic DNA (Norgen). The isolation was performed according to the manufacturer’s instructions. Prior to cDNA generation, samples were measured with the Qubit PicoGreen Assay (to assess RNA quality). Samples were then processed with the SmartSeq2 protocol (Picelli et al., 2014). Prior to sequencing, library size distribution and concentration were assessed with the bioanalyzer High Sensitivity DNA assay. For all single-cell RNA-sequencing experiments, single-cell suspensions were immediately processed with the 10x genomics chromium instrument after FACS sorting. GEM formation, reverse transcription, barcoding, cDNA amplification, and purification were all done in accordance with the manufacturer’s recommendations. Both v2 and v3 versions of the kit (10x Genomics, single cell RNA-sequencing 3’) were used. Single-cell libraries were sequenced on the NextSeq at 75 cycles with a paired end protocol. Single-cell samples were sequenced to an average read depth of 264 million reads per sample with a median count of 3 376 UMIs per cell. Bulk RNA-sequenced samples had a read depth of 15 to 20 million reads. The AP.Adamts2.1-GFP and AP.Baz1a.1-GFP samples were sequenced on the HiSeq 2500 (50 bp, single-end), while the AP.Agmat.1-GFP samples were sequenced on the NovaSeq (2x 50 bp, paired-end). The change in sequencing instrument and protocol was a consequence of logistical changes made at the institute.

### Single-cell RNA-seq analysis

Initial processing, up until the construction of the cell-barcode count matrices, of the single-cell sequencing data was done with Cell Ranger software package. Reads were mapped against a custom genome constructed from the Genome Reference Consortium Mouse Build 39 with a version 104 GTF file. Viral expression was accounted for by amending the CaMPARI2 plasmid sequence (Addgene) to the end of the FASTA file and the GTF file was edited to include the regions present on the viral mRNA. GTF and FASTA edits were performed in python and the final genome build was constructed with Cell Ranger. All subsequent analyses on the raw cell feature barcode matrices were done in R.

Expression data was imported with the SCATER package (McCarthy et al., 2017). Cells were distinguished from empty droplets with DropletUTILS (Lun et al., 2019), using a FDR of 0.05 % and a lower limit for cell detection that was calculated on a per dataset basis or set to the median UMI count for droplets greater than 10 UMIs. Cells with a mitochondrial read percentage greater than 20 % as well as cells with read counts that were more than three standard deviations below the sample average were excluded.

Feature selection and UMI correction were done with SCtransform from the Seurat package (Hafemeister and Satija, 2019; Satija et al., 2015), initially 2000 features were selected and mitochondrial percentage, sample effects, 10x chemistry (v2 vs v3), and total feature counts were regressed out. Selected features were filtered to remove any features that were differentially expressed between groups of cells separated by read count or mitochondrial percentage, thus reducing the impact of cell quality on the analysis. The final corrected UMI matrix was then used for alignment with a reference dataset (Tasic et al., 2018).

To align cells from this study with the reference dataset (Tasic et al., 2018), excluding L5 and L6 neurons, we used the Liger package (Welch et al., 2019) with a k-value of 12 and lambda-value of 10. These parameters were chosen based on how well groups were separated on a UMAP plot, as well as the expression of various markers normally enriched in each group (**Figure S2**). Then, iNMF factors were used to assign each cell from this study to a group defined in the reference dataset (L4, L2/3, SST etc.) using a weighted nearest neighbor algorithm, where the largest group sum for the of the inverse of the distance for the 10 closest reference cells was used to assign group identity. Cells whose distance to the closest cell in the reference dataset exceeded 0.002 in the iNMF factor space were removed from the dataset. Additionally, we excluded cell types that contained less than 500 cells across the entire dataset. Furthermore, the initially assigned *Pvalb* group contained a large number of cells expressing excitatory neuron markers, graphical clustering was used to subset this group and isolate *Pvalb* positive neurons that were negative for excitatory markers. L2/3 subtypes (*Adamts2, Rrad*, and *Agmat*) were determined separately within the L2/3 group with scMAP (Kiselev et al., 2018) using the scmapCluster function; 250 features were used with no similarity cutoff. These parameters allowed a demonstrably accurate assignment of L2/3 neurons to their corresponding cell types from the reference dataset (Tasic et al., 2018) types (**Figure S2**).

To analyze how the proportion of L2/3 types changed as a function of photoconversion (**Figures 2E, 2F, 2G and 2H**), we estimated the fraction of each L2/3 type (within each experiment or condition) by pooling the number of cells collected across photoconversion conditions, correcting for differences in cell number and then calculating the proportion of each cell type in these pooled datasets. These estimates allowed us to simulate a randomized distribution of proportion values for each L2/3 cell type in each experiment. Then, experimentally obtained proportion values were compared to a distribution obtained by bootstrap sampling from the combined population across photoconversion conditions.

### Artificial promoter design

Artificial promoters for selective expression in L2/3 excitatory neuron types were designed based on the ATG proximal sequence design strategy described previously (Jüttner et al., 2019). Artificial promoters contained a sequence of 2 kb surrounding the transcription start site of the endogenous marker gene taken from the UCSC genome browser (GRCm38/mm10). For the AP.Adamts2.1 promoter, the 2 kb sequence originally contained a Kozak sequence (corresponding to the translation start site of the *Adamts2* gene). To enhance transgene expression, the endogenous start codon (‘ATG’) of the Kozak sequence was changed to a stop codon (‘TGA’).

### Virus production

Artificial promoter constructs were synthesized and subcloned into an AAV vector backbone plasmid by a commercial supplier (Genscript), placed upstream of GFP for histology and gene expression experiments, and upstream of GCaMP7f (Dana et al., 2019) (modified from Addgene plasmid #104495), or a conditional version of GCaMP8s (Zhang et al., 2021) for calcium imaging experiments.

AAV vectors were produced by the FMI vector core following standard protocols. In brief, HEK293T cells were grown to 80 % confluence and co-transfected with an AAV transgene plasmid, an AAV helper plasmid, and a plasmid containing the Rep/Cap proteins (Addgene Plasmid #112862) using polyethylenimine (Polysciences). Cells were harvested and pelleted in a benchtop centrifuge 60 h after transfection and the supernatant discarded. Cell pellets were either stored at -80 °C or immediately processed. Cell pellets were resuspended in 15 ml lysis buffer (150 mM NaCl, 20 mM Tris, pH 8.0).

Solutions went through 4 freeze-thaw cycles alternating between liquid nitrogen and a 37°C water bath (15 min each) interrupted by brief periods of mixing on a Vortex shaker. Subsequently, MgCl_2_ was added to a final concentration of 1 mM, followed by the addition of Turbonuclease (bpsBioscience) at 250 U/nl. Then samples were incubated at 37 °C for 15 min. Virus solutions were spun down and the pellet was discarded. The supernatant was transferred into a prepared iodixanol OptiPrep (Sigma) gradient (6 ml 17 %, 6 ml 25 %, 5 ml 40 % and 5 ml 60 %) contained within an OptiSeal tube (Beckman,). Gradients were spun at 242 000 g at 16 °C and the viral fraction was harvested from the 40 % fraction with a 21-gauge needle. The viral fraction was further purified with Amicon 100K columns (Merck) according to the manufacturer’s instructions. Viral aliquots were stored at -80 °C.

Titers of AAV vectors were determined by qPCR. First, extra-capsid DNA was digested with DNase, 10 U/µl (Roche) for 15 min at 37 °C, and inactivated for 10 min at 95 °C. Viral capsids were then digested with proteinase K at a concentration of 100 ng/ml (Macherey Nagel) for 15 min at 37 °C and inactivated at 95 °C for 10 min. All AAVs were measured relative to standards with primers targeted to the ITRs (forward: 5′-GGAACCCCTAGTGATGGAGTT, reverse: 5′-CGGCCTCAGTGAGCGA) with the SYBR Green PCR Master Mix (ThermoFisher). qPCR was performed on the StepOnePlus real-time PCR system. Then qPCR curves were analyzed with the Step One v2.3 software from Agilent.

### Histology

To examine the viral expression pattern for artificial promoter constructs, mice were transcardially perfused with phosphate buffered saline (PBS) until the liver was exsanguinated. Mice were then perfused with 4% paraformaldehyde (PFA) in PBS. Brains were removed and placed into cold (4 °C) PFA and fixed overnight. The following day, brains were transferred to cold PBS and washed 3 times for 20 min (4 °C). Brains were sectioned with either a vibratome (Leica VT1000 S) or a cryostat (CryoStar NX70). For cryosections, brains were cryoprotected with 3 overnight washes in 30 % sucrose in PBS. Afterwards brains were equilibrated in Tissue-Tek O.C.T. compound (Biosytems) and then flash frozen in OCT with liquid nitrogen. Brains were sectioned at 20 μm and adhered to slides (FisherScientific). For vibratome sections, brains were sliced in room temperature PBS at 50 μm, stored in a 48 well plate, and then mounted on glass slides. Sections were mounted in VectaShield HardSet with DAPI (VectorLabs). Most slices also underwent a staining protocol prior to mounting. Sections were washed 4 times in PBS with 0.1 % Triton (PBST) for 10 min. Then the sections were blocked for 2 h at room temperature in PBST with 5 % neutral goat serum. Thereafter, the sections were incubated overnight at 4 °C with chicken anti-GFP (ThermoFisher). Then sections were washed in PBST 4 times for 10 min. Afterwards, the sections were stained with secondary antibodies (1:500) along with NeuroTrace 640/660 (ThermoFisher) which was diluted (1:300), followed by a final washing step (4 times 10 min PBST). Images were acquired on a Zeiss Axio Scan.Z1 slide scanner and in some cases on an Axio Imager Z2 with a LSM 700 scan head.

### Histology analysis

To determine the expression profile of viral vectors as a function of cortical depth, we used images corresponding to 200 µm in width and spanning the entire depth of cortex. Image processing was done in FIJI where images were converted to 8-bit and then enhanced for local contrast (CLAHE plugin, blocksize = 30, histogram = 256, maximum = 3). The images were then binarized with a cutoff value corresponding to the 92.5^th^ percentile of pixel intensities (calculated for each image separately) and filtered for cellular particles. Binarized images were converted to depth profiles by averaging in the horizontal dimension. All vectors were then scaled to an equal length and normalized to peak. Finally, vectors were averaged across groups, normalized to peak, and passed through a Savitzky-Golay filter.

### Bulk RNA-seq analysis

To examine the differences in expression between neurons infected with different artificial promoters, a custom genome was constructed that excluded all DNA segments used in the artificial promoter viruses. This was done to increase the accuracy of the read counts for the marker genes. A custom GTF file was used to include the viral genomes. As with the single-cell RNA-sequencing analysis, the custom genome was based on the Genome Reference Consortium Mouse Build 39 with a version 104 GTF file. Differences in sequencing method (paired vs single-end) were taken into account for the trimming and mapping steps. Reads were trimmed with the Cutadapt package (trim sequence: CTGTCTCTTATA). Gene counts were calculated with HTSeq count. Libraries were corrected for batch effects related to sequencing runs. Count values were normalized to library depth as counts per million (cpm). Genes with less than 5 cpm on average, across datasets, were excluded. Statistics were performed on fold change values.

### Two-photon calcium imaging

Calcium imaging was confined to L2/3 of V1 and was performed as previously described (Leinweber et al., 2014). A modified Thorlabs Bergamo II microscope with a 16x, 0.8 NA objective (Nikon N16XLWD-PF) was used for imaging. The illumination source was a tunable femtosecond laser (Insight, Spectra Physics or Chameleon, Coherent) tuned to 930 nm for the GCaMP imaging experiments, to 950 nm for the functional CaMPARI2 experiments, and to 1040 nm for the simultaneous imaging of red and green channels to characterize the extent of photoconversion. Emission light was band-pass filtered using a 525/50 nm or 607/70 nm filter (Semrock), and signals were detected with a GaAsP photomultiplier (Hamamatsu, H7422). PMT signals were amplified (Femto, DHCPCA-100), digitized at 800 MHz (National Instruments, NI5772), and band-pass filtered at 80 MHz with a digital Fourier-transform filter on a field-programmable gate array (National Instruments, PXIe-7965). The microscope scan head was a 12 kHz resonant scanner (Cambridge Technology). Image acquisition was at a resolution of 750 pixels by 400 pixels (60 Hz frame rate) with a field of view of 375 μm by 300 μm. Four layers, separated by 15 μm were imaged sequentially with a piezo-electric linear actuator (Physik Instrumente, P-726), and the effective frame rate per layer was 15 Hz.

### Two-photon image analysis

Image analysis was performed as previously described (Keller et al., 2012). In brief, acquired data were full-frame registered to correct for brain motion. In datasets with weaker signal, registration was performed on a running average over 5 to 10 frames. Raw fluorescence traces for individual neurons were calculated as the average signal in a region of interest (ROI) that was manually selected based on mean and maximum fluorescence images. To correct for slow drift in fluorescence, the raw fluorescence trace was first corrected with an 8^th^ percentile filter over a window of 66 s (1000 frames) as described previously (Dombeck et al., 2007), and F_0_ was defined as the median fluorescence over the entire trace. To calculate the average response traces, we first calculated the average event-triggered fluorescence trace for each neuron. The responses of all neurons were averaged across triggers and baseline subtracted. The baseline window for the CaMPARI2 analysis was 1.33 s (20 frames) and for the GCaMP analysis it was 0.66 s (10 frames). We used different analysis windows for the two indicators due to the differences in signal to noise levels. To quantify the difference of average calcium responses, for GCaMP based analyses (**Figure 4**), as a function of time, we performed a two-sample t-test for every bin of the calcium trace (15 Hz) and marked bins as significantly different for p < 0.001. For visual clarity, we removed isolated selected bins, such that a significant bin was only marked if at least two neighboring bins were also significant.

## Key Resources Table

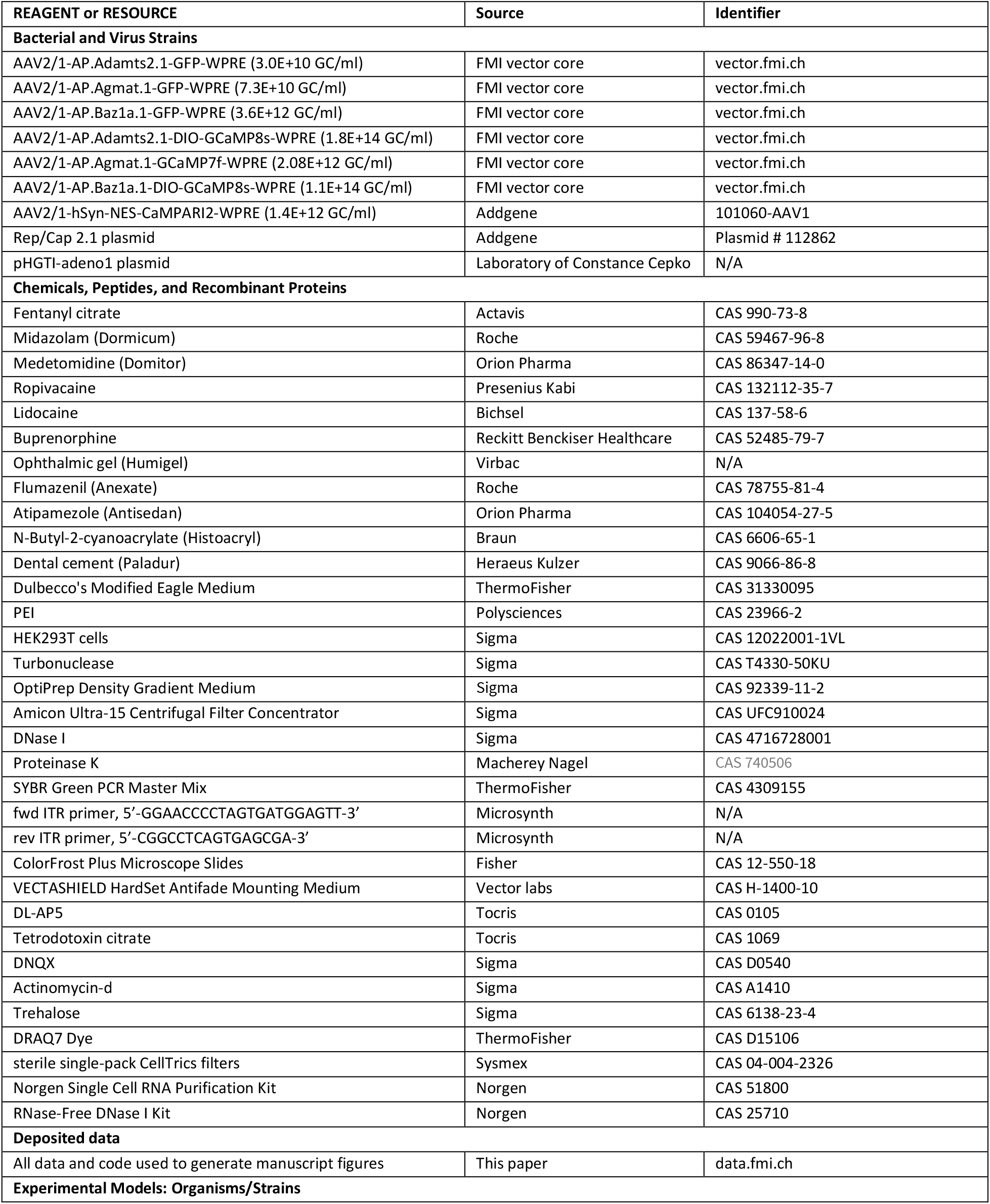
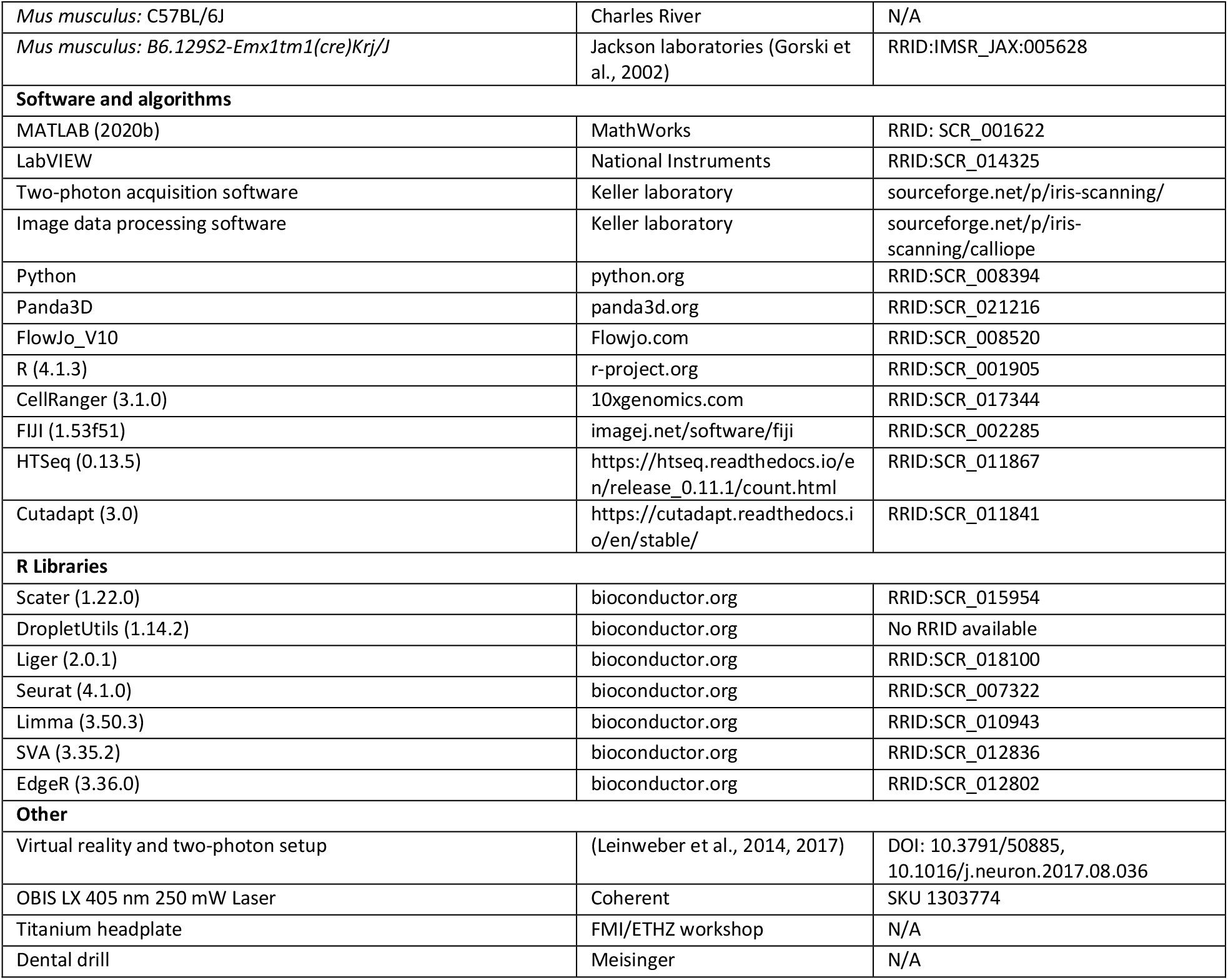

## Data and code availability

Software for controlling the two-photon microscope and preprocessing of calcium imaging data is available on https://sourceforge.net/projects/iris-scanning/. Raw data and code to generate all figures of this manuscript are available on https://data.fmi.ch/PublicationSupplementRepo/.

**Table S1.**
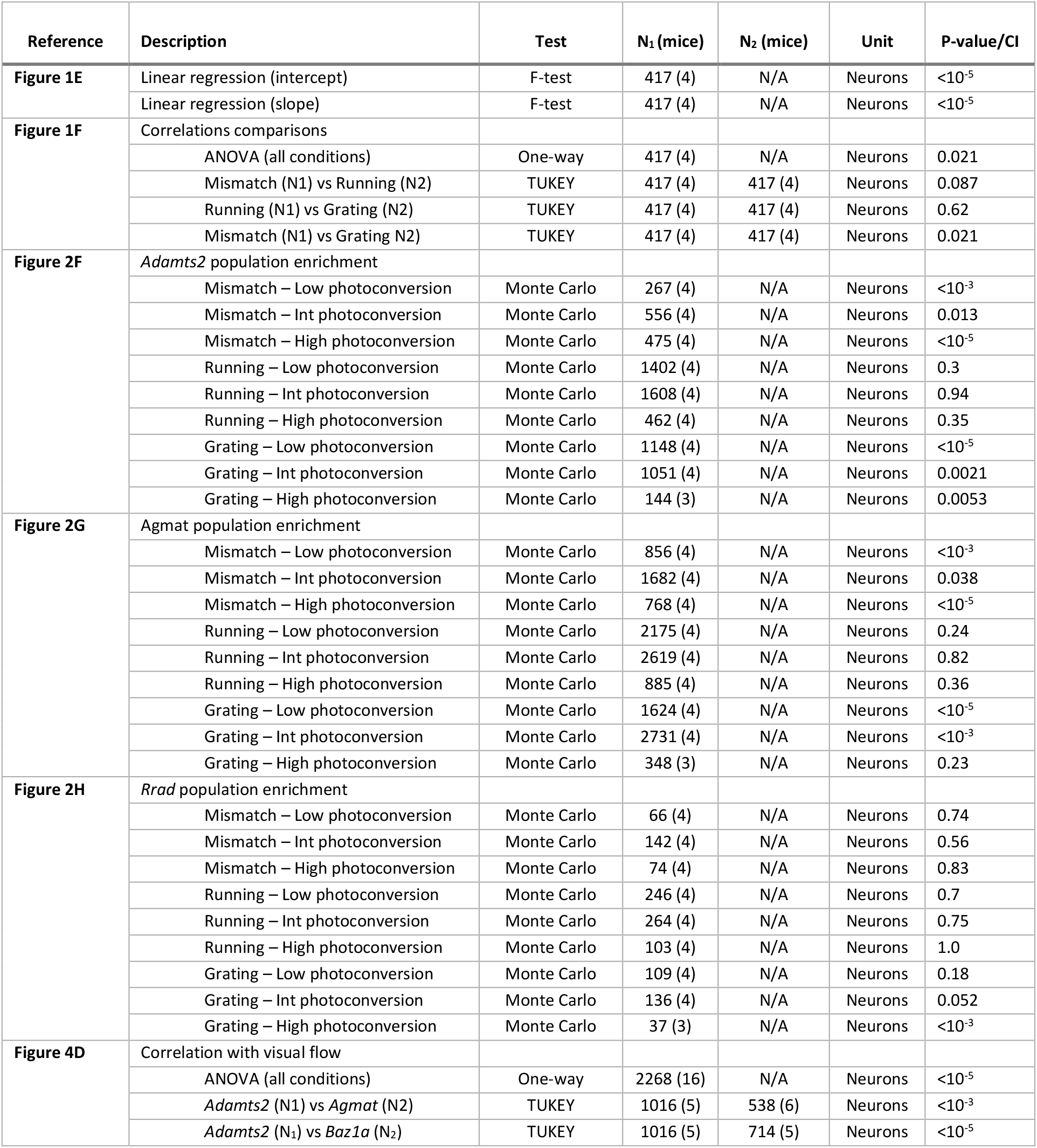
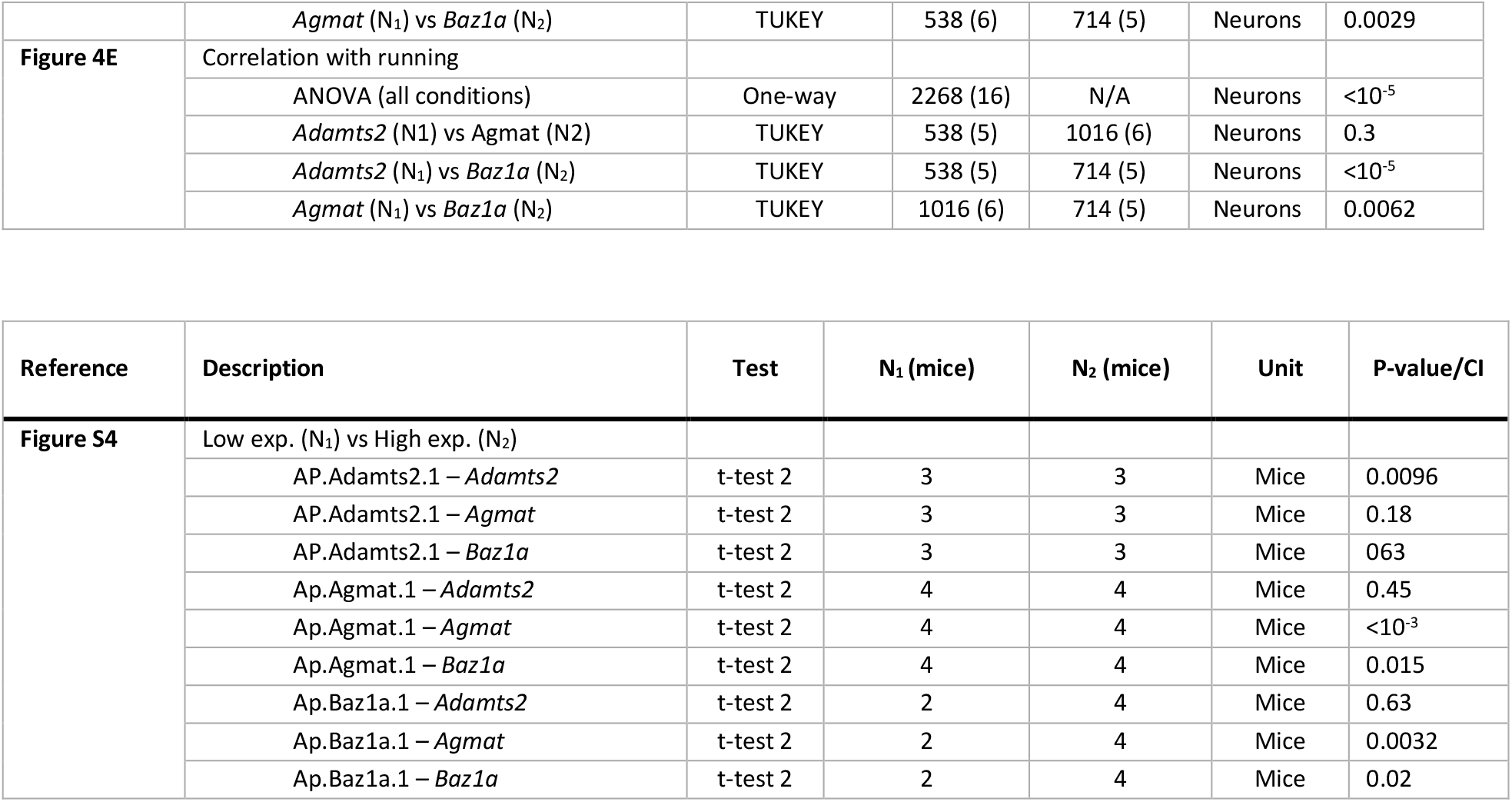
Statistics. All values are rounded to two significant decimals, except values smaller than 10^−3^. The tests used, were two-sample independent t-test (t-test 2), one way ANOVA with a post-hoc TUKEY, and Monte Carlo simulations. Sample sizes were estimated based on previous experiments performed in the same behavioral paradigms (Attinger et al., 2017; Heindorf et al., 2018).

**Table S2.** Number of mice per experiment. The table below lists the number of mice in each experiment.

| Dataset | Number of mice |
| --- | --- |
| Mismatch-photoconversion, functional imaging (Figures 1D - 1F) | 4 |
| Mismatch-photoconversion, single-cell (Figures 2, S2) | 4 |
| Running-photoconversion, single-cell (Figures 2, S2) | 4 |
| Gratings-photoconversion, single-cell (Figures 2, S2) | 4 |
| AP.Adamts2.1-GFP, histology (Figures 3B and 3F) | 4 |
| AP.Agmat.1-GFP, histology (Figures 3C, 3E and 3F) | 2 |
| AP.Baz1a.1-GFP, histology (Figures 3D and 3F) | 2 |
| AP.Adamts2.1-GFP, bulk RNA-seq (Figures S4A, S4D and S4G) | 3 |
| AP.Agmat.1-GFP, bulk RNA-seq (Figures S4B, S4E and S4H) | 4 |
| AP.Baz1a.1-GFP, bulk RNA-seq (Figures S4C, S4F and S4I) | 4 |
| AP.Adamts2.1-GCaMP (Figures 4A - 4E, S5) | 5 |
| AP.Agmat.1-GCaMP (Figures 4A - 4E, S5) | 6 |
| AP.Baz1a.1-GCaMP (Figures 4A - 4E, S5) | 5 |
| Ef1α-GCaMP (Figures S5A and S5B) | 16 |

## Acknowledgements

We thank all the members of the Keller lab for discussion and support as well as Tingjia Lu and Daniela Gerosa for vector production. We thank Sirisha Aluri, Sebastien Smallwood and Hubertus Kohler for help with the single-cell sequencing experiments as well as Koshika Yadava for help with the histology. This project has received funding from the Swiss National Science Foundation, the Novartis Research Foundation, and the European Research Council (ERC) under the European Union’s Horizon 2020 research and innovation programme (grant agreement No. 865617).

## Author contributions

SO designed and performed the experiments and analyzed the data. HO designed and performed experiments. S.O. and G.K. wrote the manuscript.

